# A sequence- and structure-based characterization of microbial enzymes identifies P. stutzeri as a plastic-degrading species

**DOI:** 10.1101/2024.04.12.589142

**Authors:** Alexander Hong, Serafina Turner, Rita Glazer, Zachary A. Weishampel, Atharva Vispute, Ashley Huang, Zachary A. Holmes, Beatrice Schleupner, Meagan M. Dunphy-Daly, William C. Eward, Jason A. Somarelli

## Abstract

Plastic waste has accumulated rapidly in the past century and is now found throughout every ecosystem on Earth. Its ubiquitous presence means that plastic is routinely ingested by countless organisms, with potential negative consequences for organismal health. New solutions are urgently needed to combat plastic pollution. Among the many strategies required to curb the plastic pollution crisis, the bioremediation of plastic via enzymatic activity of microbial species represents a promising approach. Diverse microbes harbor enzymes capable of degrading plastic polymers and utilizing the polymers as a carbon source. Herein, we characterize the landscape of microbial protein-coding sequences with potential plastic degrading capability. Using the two enzyme systems of PETase and MHETase as a guide, we combined sequence motif analysis, phylogenetic inference, and machine learning-guided 3D protein structure prediction to pinpoint potential plastic-degrading enzymes. Our analysis platform identified hundreds of enzymes from diverse microbial taxa with similarity to known PETases, and far fewer enzymes with similarity to known MHETases. Phylogenetic reconstruction revealed that the plastic degrading enzymes formed distinct clades from the sequences of ancestral enzymes. Among the potential candidate sequences, we pinpointed both a PETase-like and MHETase-like enzyme within the bacterium *Pseudomonas stutzeri*. Using plate clearing assays, we demonstrated that *P. stutzeri* is capable of degrading both polyurethane (Impranil®) and polycaprolactone (PCL). *Pseudomonas stutzeri* also grew on carbon-free agar supplemented with polystyrene, suggesting this organism can utilize synthetic polymers as a carbon source. Overall, our integrated bioinformatics and experimental approach provides a rapid and low-cost solution to identify and test novel polymer-degrading enzymes for use in the development of plastic bioremediation technologies.

## Introduction

The advent of synthetic plastic polymers has dramatically reshaped nearly all facets of our daily lives. From construction materials to food packaging (Lebreton & Andrady, 2019), plastic pervades our modern-day existence. Initially lauded for its durability and diverse applications, these advantageous features of plastic bring unintended negative consequences as plastic waste continues to rapidly accumulate globally. Every year, between 9 and 23 million metric tons of plastic waste end up in rivers, lakes, and oceans (MacLeod et al., 2021). By 2050, it is predicted that there will be more plastic debris in the ocean than fish (World Economic Forum Industry Agenda, 2016). Plastic pollution is a global threat that continues to pose potential risks to the environment and organismal health (Morrison et al., 2022) as organisms can become entangled in waste or unintentionally ingest microplastics (Webb et al., 2012). Additionally, there is growing concern over the unknown consequences of chemical additives that may leach from plastics into our water and food supplies, may adsorb harmful chemical compounds, and may impact well-being over a lifetime of exposure (Cherif Lahimer et al., 2017).

Current methods of disposing of plastic products, such as chemical degradation and incineration, are neither environmentally friendly nor sustainable. These processes require high pressure and temperature conditions that are expensive, energy-consuming, and inefficient (Webb et al., 2012). Disposal methods of this kind also lead to the generation of harmful waste products, including polycyclic aromatic hydrocarbons, polychlorinated biphenyls (PCBs), and heavy metals, which are associated with adverse health effects (Webb et al., 2012). Likewise, the effectiveness of mechanical and chemical recycling is limited by additives and impurities in plastic (Schyns & Shaver, 2021).

Novel bioremediation strategies are another approach to address the plastic waste. Bioremediation is a process in which organisms, either genetically-engineered or naturally-occurring, are employed to metabolize and thereby remediate environmental contaminants. Bioremediation has been used to decontaminate sites laden with heavy metals, oil, and radioactive materials (Abo-Alkasem et al., 2023). These approaches leverage microorganisms or plants to convert toxic substances into non-toxic byproducts or to sequester the contaminants in biologic material for later remediation or containment (Prakash et al., 2013). To date, numerous plastic degrading species have been identified, including bacteria, fungi, insects, and crustaceans (Dawson et al., 2018); (Sheth et al., 2019).

Much of the focus to date has been on the biotransformation of polyethylene terephthalate (PET) by naturally-evolved PET hydrolases. Multiple PET hydrolases have been identified, many of which evolved from cutinases, a family of serine esterases that can catalyze reactions with triacylglycerols and polyesters (Yoshida et al., 2021). One of the first reported PET hydrolases was a poly(butylene terephthalate-co-adipate) (BTA)-hydrolase from *Thermobifida fusca*. Since then, multiple enzymes have been identified with the ability to degrade PET, notably the PETase in *Ideonella sakaiensis*, Cut190 in *Saccharomonospora viridis*, and LC-cutinase from leaf compost (Kawai, 2021; Sulaiman et al., 2012; Yoshida et al., 2021). These discoveries of naturally-evolved enzymes have paved the way for bioengineering and machine learning-based strategies to improve on these natural systems to enhance rates of PET degradation (Lu et al., 2022; Richter et al., 2023; Tournier et al., 2020). Naturally-evolved enzymes provide a template for bioengineers to develop more specific and efficient enzymes.

The utility of naturally-evolved enzymes in bioengineering highlights the importance of understanding the landscape of naturally-evolved plastic-degrading enzymes. To this end, we employed a sequence- and structure-based analysis to identify candidate plastic degrading enzymes across diverse microbial taxa. Using a group of known PETase and MHETase enzymes as reference inputs, we uncovered thousands of sequences with high sequence similarity to known PETases and hundreds with similarity to known MHETases. Phylogenetic inference further highlighted subsets of PETase-like and MHETase-like enzymes that form clades with known PETases and MHETases. Cross-checking the species annotations for these candidate enzymes, we identified *Pseudomonas stutzeri* as possessing both a PETase-like and MHETase-like enzyme. We used AlphaFold2.0 to predict the 3D structures for the *P. stutzeri* enzymes. The inferred structures resemble those of known bacterial plastic-degrading enzymes. We validated the plastic degrading capacity of *P. stutzeri* by using plate clearing assays (PCAs) with both Impranil® DLN-SD and polycaprolactone (PCL). Growth on carbon-free agar enriched with polystyrene suggested that *P. stutzeri* may be capable of utilizing synthetic polymers as a carbon source. Our rapid and low-cost analysis platform enabled the identification and prioritization of candidate species for subsequent experimental testing. Notably, this approach led to the discovery of *P. stutzeri* as a novel plastic-degrading bacterial species.

## Methods

### Identification and characterization of potential plastic-degrading enzyme sequences

To identify new bacterial PETase-like and MHETase-like enzymes, nine known bacterial PETases, three known fungal PETases, and three known MHETases were used to query the National Center for Biotechnology Information (NCBI) sequence database using the protein basic local alignment and search tool (BLASTp) with low stringency filters (expect threshold = 20, gap existence = 9, extension = 1). A list of sequences in FASTA format was compiled for each query (**Supplemental Table 1**).

Known PETase sequence motifs (oxyanion hole, catalytic triad, GXSMGGGG motif (Danso et al., 2018; Dimitriou et al., 2019)), were identified using multiple sequence alignments. Additional sequence motifs and active sites were identified from multiple sequence alignments (DECIPHER package in R Studio) (E. Wright, 2017) using the MEME Suite (Bailey et al., 2015) in discriminative mode. An outgroup for MEME was constructed from the closest protein family of the query category obtained from a batch search of the NCBI Conserved Domain Database (CDD). The motifs and active sites common to each group identified using MEME were used to filter the list of proteins generated in the low-stringency BLAST queries. The resulting lists of PETase- and MHETase-like enzymes were compared for organism overlap (**Figure 1A**).

**Figure 1:**
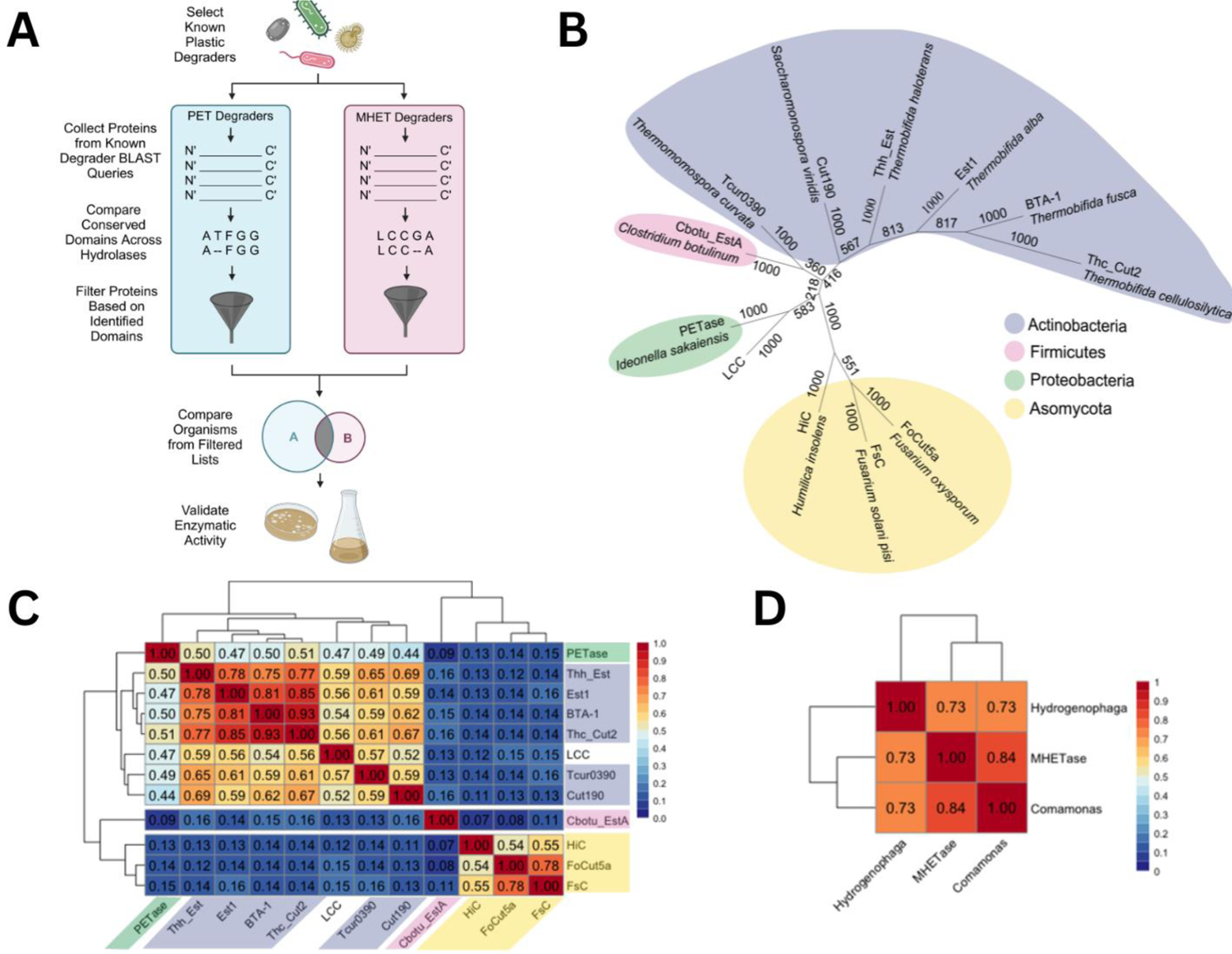
Phylogenetic analysis of PET and MHET hydrolases reveals similarity between known plastic degraders. A. Schematic of the analysis platform, including comparison of species for overlap in those containing both PETase-like and MHETase-like sequences for experimental validation. B. Unrooted maximum likelihood phylogenetic tree of the nine bacterial and three fungal species with experimentally proven PET hydrolases used in this study, color coded by phylum. Phylum for LCC is unknown as it comes from uncultured bacteria. C. Percent identity heat map of the experimentally validated PET hydrolases in this study. D. Percent identity heat map of the three MHET hydrolases. Red indicates a high percent identity, while blue indicates a low percent identity.

### Phylogenetic analysis of PET and MHET degraders

Multiple sequence alignments were generated using the DECIPHER software. Phylogenetic trees were constructed using maximum likelihood with 1,000 bootstrap replicates using PHYLIP (Felsenstein, 1989). Maximum likelihood algorithms infer the probability of observing a sequence, given a specific phylogenetic tree structure.

### Structural analysis of PET and MHET degraders

All 3D protein structures were predicted using the deep-learning artificial intelligence system, AlphaFold2.0. This was executed on the Duke Compute Cluster (DCC) using a job script and a FASTA file containing the amino acid sequence as input. The “ranked_0.pdb” structure, representing the most confident prediction, was selected for further analysis (Jumper et al., 2021). Predicted 3D structures were visualized and superimposed in PyMOL (version 2.5.2, Schrödinger, LLC., 2021) to calculate root-mean-square deviation (RMSD) between pairwise superpositions. Alignments with outlier rejection (5 cycles, 2.0 cutoff) were performed to focus on the conserved folded regions, thus reducing the impact of less confident long loop regions. RMSD values less than 2.0 Å were considered good superpositions, with values less than 1.0 Å indicating excellent overlap between structures (Castro-Alvarez et al., 2017).

### Bacterial cultures

Agar plates were prepared with standard Luria-Bertani (LB) broth (10 g of tryptone, 5 g of yeast extract, 10 g of NaCl, per 1 L of deionized sterile water) for the strain of interest, and plates were inoculated by streaking to obtain single colonies. Plates were incubated overnight at 37°C in a humidified incubator and single colonies were used to inoculate 10 mL liquid broth cultures. Cultures were incubated at 37°C for up to 72 hours, depending on the growth rate of each bacterial isolate, until the bacteria reached log phase of growth (OD600∼1.0). Strains of *Pseudomonas (P. stutzeri)* included three *P. stutzeri* isolates ATCC 51152, ATCC 17832, and ATCC 31258. *E. coli DH5α* was included as a negative control for PCAs and was cultured using the conditions described above.

### Impranil® Plate Clearing Assay

LB agar was prepared using 10 g/L tryptone (peptone from casein), 5 g/L yeast extract, 10 g/L NaCl, and 15 g/L agar–agar solubilized in deionized water (Molitor et al., 2020). Media was autoclaved (120°C, 20 minutes) and immediately moved to a hot plate with rapid mixing with a magnetic stir bar. Agar was maintained between 40 and 80°C. A total of 4 mL of Impranil® DLN-SD emulsion (COVESTRO, Leverkusen, Germany), an anionic aliphatic polyester polyurethane solution, was added to 1 L of sterile molten LB agar, after which 15-25 mL portions of mixture were poured into 10 cm bacteriological Petri dishes, and the agar was allowed to solidify at room temperature for at least 15 minutes or until firm and transferred to a cold room.

To perform PCAs, bacteria in liquid culture were spotted on plates with an inoculating loop and incubated at optimal growth temperature in a sealed plastic container lined with paper towels soaked in sterile water. Plates were stored for a minimum of three days and up to seven days as this was experimentally determined to be the best incubation time for *P. stutzeri*. If plate clearing had not occurred during this timeframe, plates were moved to the 4°C cold room to reduce rate of growth in an attempt to allow the colony to produce more enzymes. Positive clones were identified by a visible halo of cleared agar-polymer mixture. Agar plates were imaged with a digital camera or in a light box (Oxford Optronix GelCount, UK).

### PCL Plate Clearing Assay

To create PCL agar plates, the polymer was emulsified in LB agar. 0.1 g of PCL granules (average Mn °10,000 by GPC, density 1.146 g/mL, Sigma-Aldrich/Merck) were added per 20 mL of acetone in a 50 mL Erlenmeyer flask. The mixture was heated to 50°C and stirred at 500 RPM using a magnetic stir bar in a fume hood for approximately 15 minutes, at which point the liquid became completely clear. The PCL emulsion was then added drop by drop into the liquid agar, resulting in a slightly translucent color. The mixture was then heated to 90°C to boil off the acetone until the volume reduced to the original liquid agar volume. Mixtures were poured into Petri dishes, then stored at 4°C in a sealed bag to prevent agar dehydration. To screen bacteria for plate clearing, plates were inoculated again with an inoculation loop and placed upside down in a 37°C incubator inside a plate humidifier. Plates were checked daily for clearing and if no plate clearing occurred, the plates were then moved to the cold room similar to the Impranil*®* plate clearing assay.

### Carbon-Free Polystyrene Growth Assay

Fresh bacterial cultures were prepared from glycerol stocks. To create the LB plates, an agar mix of LB base was created and autoclaved and 30 mL was poured into 10 cm bacteriological Petri dishes. Carbon-free Bushnell Haas (BH) agar plates were made using 20 mL of a BH base layer and 15 mL of a top layer of BH agar infused with 9.5-11.5 µm polystyrene spheres (Cospheric, Santa Barbara, CA). The top layer was created by mixing 500 mg of each microsphere size into 50 mL of the BH agar mixture. This mixture was then autoclaved and 15 mL of the polymer agar was poured into the already-set 20 mL BH plates. Prior to testing, plates were stored at 4°C in a sealed bag to prevent dehydration. To test bacteria for utilization of polystyrene, plates were spotted in three distinct regions with 0.6 μL of bacteria and placed upside down in a 37°C incubator inside a plate humidifier. Plates were monitored daily for growth.

## Results

### Known plastic degrading enzymes from diverse taxa share sequence and structural similarities

We characterized a suite of PETase-like enzymes across diverse microbial taxa. To do this, we developed an analysis workflow using known plastic-degrading enzymes as input queries to search publicly-available microbial genomes (**Figure 1A**). Phylogenetic analysis of the input queries revealed four clades with robust bootstrap support (**Figure 1B**). The clades were composed of 1) bacterial cutinases and esterases, such as Cut190 (Kawai et al., 2020) and others; 2) the *C. botulinum* esterase, EstA (Perz et al., 2016); 3) *I. sakaiensis* PETase and a leaf branch cutinase with sequence similarity to *T. fusca* (Sulaiman et al., 2012); and 4) fungal PETases (**Figure 1B**). Using sequence alignment, we found high sequence similarity among bacterial queries, but low similarity between EstA and other bacterial sequences. Similarly, fungal PETases had high sequence conservation, but differed substantially from bacterial PETases (**Figure 1C**). These results suggest PETase-like enzymes may have evolved through multiple, independent evolutionary pathways. In a parallel approach, we also used known MHETases - which had 70% sequence identity (**Figure 1D**) - as queries to search NCBI.

Using AlphaFold2.0, we predicted the 3D structures of the known bacterial PETases and superimposed these structures using PyMOL. All PETase-like enzymes had highly similar structures, as indicated by their close superposition (RMSD values close to 1 indicate near perfect superposition). (**Figure 2A**). It is noteworthy that although the sequence similarity of EstA from *C. botulinum* is relatively low compared with other PETase-like enzymes, the 3D structure overlaps tightly with other PETase-like enzymes (**Figure 2A**). We next used MEME to identify potential functional motifs in known bacterial MHETases. This analysis suggested three conserved motifs within the known bacterial MHETases, two of which flank the lid binding site, which is important for substrate specificity (**Figure 2B; Supplemental Figure 2**). Superposition of these known MHETases revealed close structural overlap, with RMSD values <1Å (**Figure 2C**).

**Figure 2:**
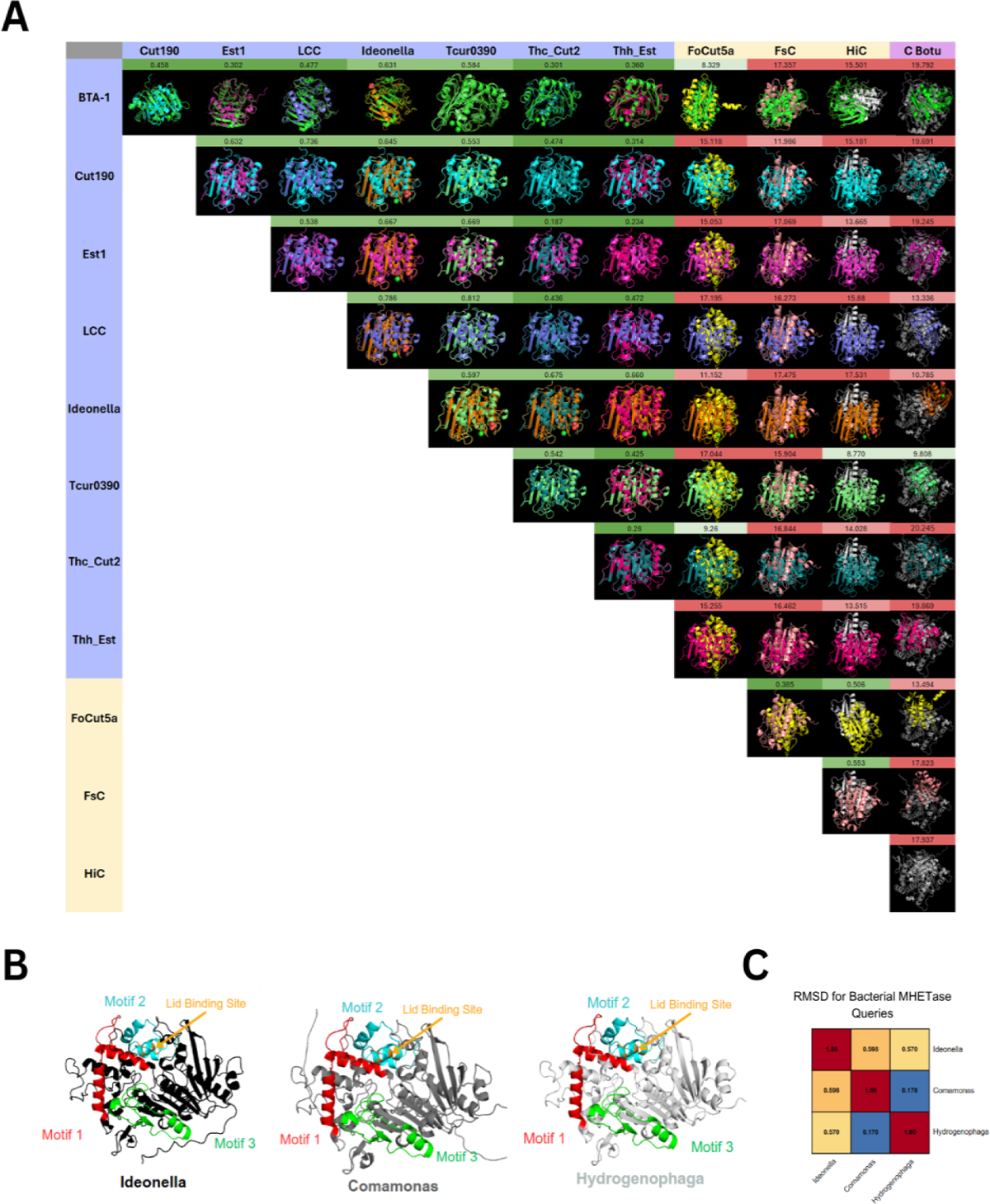
Comparison of predicted structures for known PETases and MHETases. **A.** Superposition of bacterial PET hydrolase queries. Numbers above indicate root-mean-square deviation (RMSD) of the overlap between structures. B. Predicted 3D structures of known bacterial MHETases. **C.** RMSD for superposition of known bacterial MHETases.

### Comparison of known plastic degrading enzymes with ancestral outgroups

Bacterial PETases are similar to alpha/beta hydrolases, a broad class of hydrolases with wide-ranging substrate specificity characterized by an α/β/α structure of α-helices flanking a central region of 5-8 β-sheets (Renault et al., 2005). To understand how known PETases relate to other ancestral sequences, we reconstructed phylogenetic relationships between a representative group of alpha-beta hydrolases within PFam (PF20408) and known PETases. These analyses suggest that known PETases group into a distinct clade from other alpha-beta hydrolases (**Figure 3A**). A similar trend was observed for bacterial MHETases: known MHETases formed a separate clade from other enzymes in the same protein family (PFam 07519) (**Figure 3B**).

**Figure 3:**
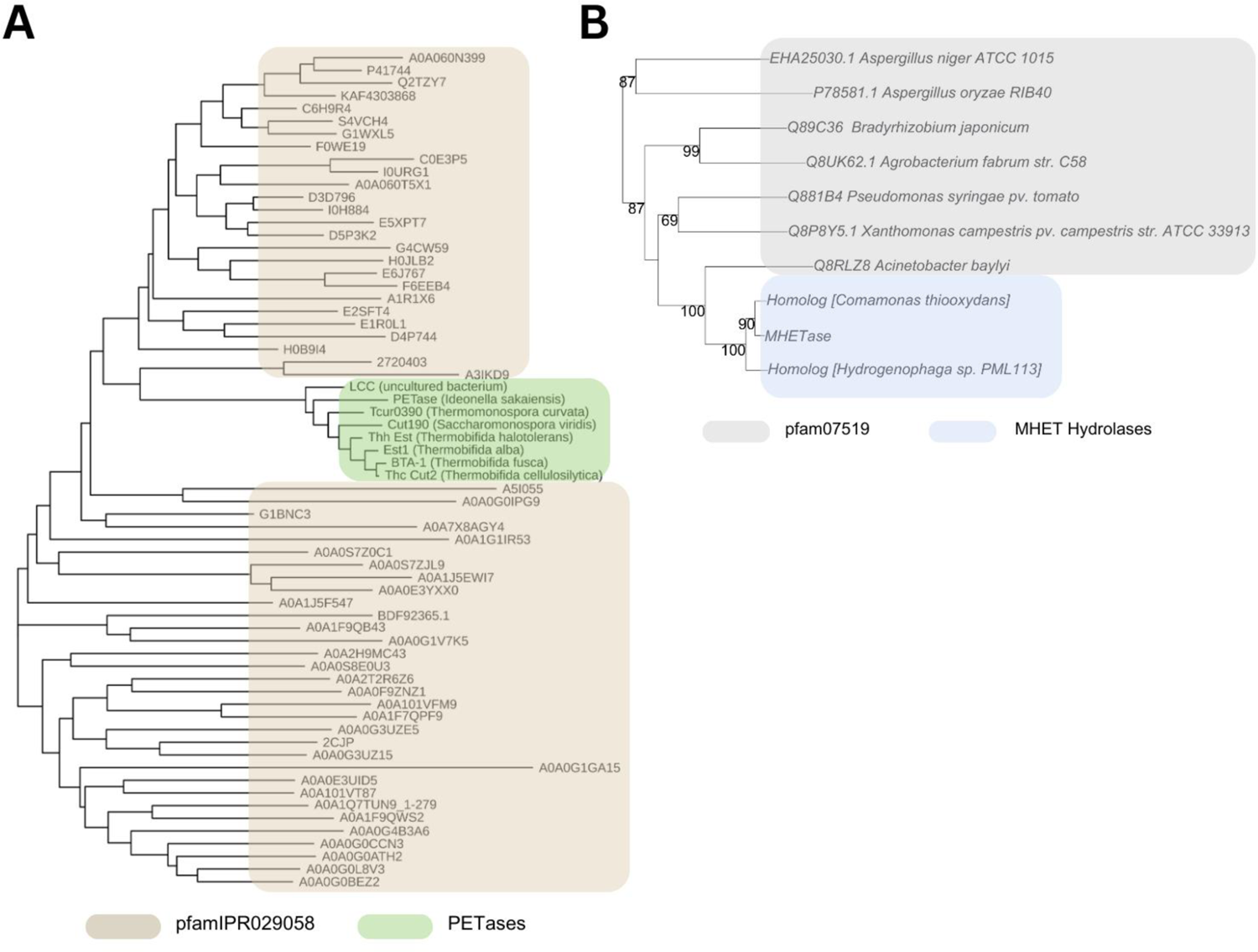
Phylogeny of PET and MHET hydrolases reveal structural overlap. **A.** Maximum Likelihood Phylogenetic Tree with PETases, alpha-beta hydrolases, and select cutinases across domain model pfamIPR029058 from ELIXIR database InterPro. Notably, the clade of PETases exhibits distinct phylogenetic divergence from other enzyme groups within the tree. **B.** Maximum Likelihood Tree of MHETase, MHETase homologs, and domain model pfam07519 in NCBI’s Conserved Domain database with corresponding bootstrap values. Pfam07519 was identified as a nonspecific hit to the three MHET hydrolases by BATCH search in the CDD. Proteins are colored by group.

Interestingly, the chlorogenate esterase from *Acinetobacter baylyi* groups most closely within the group of known MHETases, highlighting this species as a candidate for further testing of MHETase activity.

Phylogenetic reconstruction of fungal PETases with other related enzymes (PFAM01083; predominantly cutinases and esterases) revealed a clade of known PETases grouping with three enzymes: two cutinases from *Aspergillus oryzae* ((Liu et al., 2009) and *Pyricularia oryzae* (sequence ID: XM_003719278.1), and a carbohydrate esterase from *Fusarium vanettenii* (Xie et al., 2022) (**Figure 4A**).

**Figure 4:**
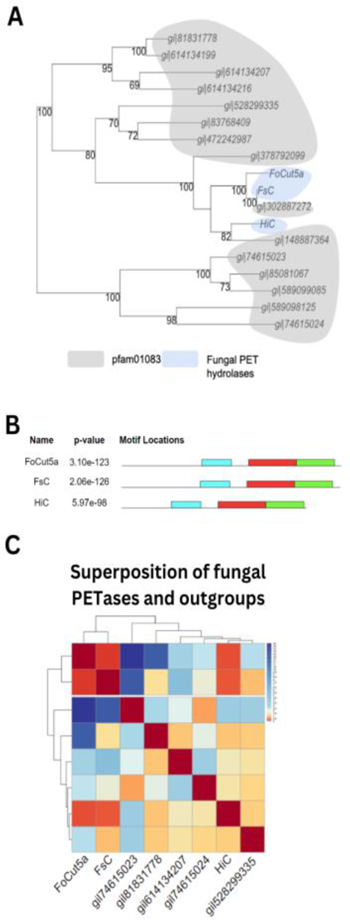
Fungal PETase analysis indicates high sequence similarity and structural overlap. **A.** Maximum Likelihood Tree of fungal PET hydrolases and cutinase domain model, pfam01083, in NCBI’s Conserved Domain database with corresponding bootstrap values. The cutinase domain was identified for all three PET hydrolases by BATCH search in the CDD. **B.** Relative location of motifs in each PET hydrolase identified by motif discovery software MEME under discriminative mode with the enzymes in pfam01083’s domain model an outgroup and PET hydrolases as an in group. **C.** Heatmap of RMSD values for each fungal PET hydrolase predicted structure (red: high similarity, blue: low similarity).

Using tBLASTn we cross-checked these species for PETase-like enzymes and identified protein-coding sequences from all three species. Interestingly, we identified a PETase-like enzyme in *Fusarium vanettenii* with 99% identity to the known fungal PETase in *Fusarium solani*. These species form a group known as the *Fusarium solani* species complex (Coleman et al., 2009). These analyses also identified a PETase-like enzyme in *Pyricularia oryzae* with 60% identity to the known PETase from *Humicola insolens* (sequence ID: XM_003719278.1) and a PETase-like enzyme in *Aspergillus oryzae* with 54% identity to the *Humicola insolens* PETase (sequence ID: XM_001825747.1).

Analysis of conserved sequences among known fungal PETases using MEME pinpointed three conserved motifs (**Figure 4B**), including a hydrophobic 29 amino acid motif in the central portion of each protein, as well as two C-terminal motifs that correspond to the alpha-beta hydrolase domain (**Figure 4B**; **Supplemental Figure 3B**).

### Identification of P. stutzeri as a novel plastic degrading species

Using bacterial hydrolases as queries, we identified a total of 33,601 sequences, of which 2,832 sequences had ≥50% identity to the query. Of the fungal protein searches, we identified 12,196 sequences, of which 558 showed ≥50% identity to their respective query sequence. For PETases, the queries produced 33,601 hits, of which 2,832 showed ≥50% identity. Using MHETases as queries produced 18,855 hits, of which 107 showed ≥50% identity (**Supplemental Table 1**).

Filtering sequences by the presence of identified motifs resulted in 1,369 bacterial species with a PETase-like enzyme and three bacterial species with a MHETase-like enzyme (**Figure 5A**). Among these species, only one species - *Pseudomonas stutzeri -* contained both a PETase-like enzyme and a MHETase-like enzyme (**Figure 5A**). *P. stutzeri* contains four PETase-like sequences: EQM75115.1, PNG04480.1, WP_102852227.1, WP_146031689.1, and modeling the 3D structure of these sequences against the *I. sakaiensis* PETase showed folds with high similarity and RMSD values of 0.506, 0.535, 0.495, and 0.541 Å, respectively (**Figure 5B**). *P. stutzeri* also contains two accession numbers of MHETase-like enzymes: NIU62236.1 and NIU62237.1. When superimposed with the *I. sakaiensis* MHET hydrolase, these exhibit relatively low RMSD values of 1.095 and 0.425 Å, respectively (**Figure 5C**).

**Figure 5.**
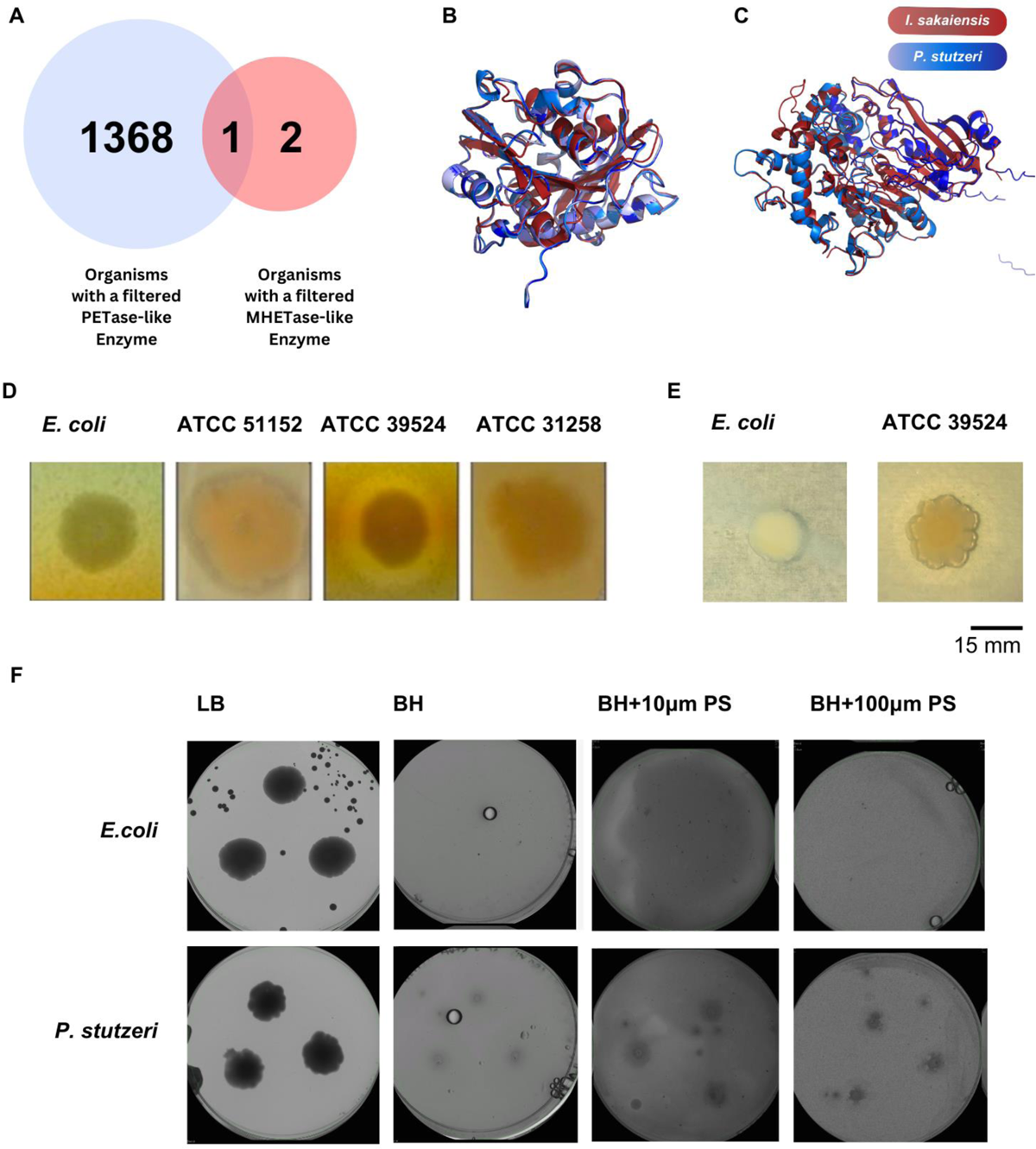
Validation of *P. stutzeri* plastic degradation and utilization. **A.** The organisms of filtered proteins of PETase-like and MHETase-like enzymes were compared for overlap. The single species with both enzymes is *Pseudomonas stutzeri.* **B.** Predicted structures of PET hydrolases and **C.** MHET hydrolases from *I. sakaiensis* (red) superimposed onto the *P. stutzeri* structures (blue). **D.** *E. coli* (negative control) and three strains of *P. stutzeri* were screened for polyesterase activity in Impranil® PCAs. **E.** Plate clearing assay results for *E. coli* and *P. stutzeri* (ATCC 17832) on PCL plates.

### Experimental validation of P. stutzeri as a novel plastic degrading organism

While *E. coli* had no visible signs of clearing, all three strains of *P. stutzeri* produced visible halos on Impranil® (**Figure 5D**). Interestingly, the different strains displayed variation in the rate of clearing. ATCC 17832 produced clearing on Impranil® DLN agar plates within two days, while ATCC 51152 and ATCC 31258 both formed discolorations around the colony after two weeks at 37°C. We also observed continued plate clearing at 4°C. ATCC 17832 was able to clear the entire plate following two weeks at 4°C, while ATCC 51152 and ATCC 31258 took approximately one month to complete the plate clearing (not shown). We further validated the ability of *P. stutzeri* to degrade plastic using plates containing PCL, a synthetic, semi-crystalline, biodegradable polyester. Consistent with the results from the Impranil® plates, *P. stutzeri* generated visible halos on PCL plates (**Figure 5E**). We further observed ATCC 17832 growth on the two BH + agar + polystyrene plates but none on the BH + agar plates, suggesting that the bacteria utilized polystyrene as a carbon source (**Figure 5F**). Together, these results suggest that *P. stutzeri* represents a novel plastic degrading bacterial species.

## Discussion

Microorganisms exhibit a remarkable capacity for adaptation to novel environments and resources. Microbial life has been discovered in diverse habitats on Earth, living in extreme temperatures (D’Amico et al., 2006; Thakur et al., 2022), salinity and radiation (Thakur et al., 2022), and even in space (Brandt et al., 2015). Microbes have adapted pathways to use diverse resources as nutrients ranging from light (photosynthesis) to chemical processes by oxidizing both inorganic (hydrogen, iron(II), sulfur, ammonium) and organic compounds (fatty acids, alcohols, alkanes, macromolecules) (Lever et al., 2015).

In addition to their use of naturally occurring nutrients and materials, a growing number of reports have also identified microbes capable of degrading different types of plastic (Sheth et al., 2019). These organisms span diverse microbial taxonomic groups, including both bacterial and fungal species (Sheth et al., 2019). Here, we used a combination of bioinformatics analysis, machine learning-based 3D protein structure prediction, and experimental validation to characterize the landscape of plastic-degrading enzymes from available bacterial and fungal genomes. Filtering species that contain both a PETase-like and MHETase-like enzyme, we identified *Pseudomonas stutzeri* as a top candidate with plastic degrading capacity and validated the ability of *P. stutzeri* to degrade plastic and utilize polystyrene as a carbon source.

Our results and those of others (Joo et al., 2018; Nelson & Ramos, 2023; Son et al., 2020) reveal thousands of potential PETase-like protein sequences spanning numerous microbial taxa. The diversity in plastic degrading species suggests multiple independent evolutionary origins for these plastic degraders. Phylogenetic reconstruction of known PETases and an outgroup of alpha/beta hydrolases suggests that these PETase-like proteins form distinct clades. These clades originate from ancestral alpha/beta hydrolases that have pre-existing capacity as esterases (Renault et al., 2005). These results suggest that the sequence differences between fungal PETases and related enzymes, such as alpha-beta hydrolases, may result in 3D structures that are unique to the fungal PETases.

Conversely, while PETase-like enzymes are more common, we identified far fewer MHETase-like sequences. Our phylogenetic analysis of MHETase-like enzymes suggests that -as with the case PETases - known MHETases form a distinct clade from ancestral tannases. This indicates there are specific mutations within experimentally-verified MHETases that distinguish them from the broader class of tannases (**Figure 2**) (Knott et al., 2020). We do not know why these MHETase-like enzymes are less frequently observed; it is possible that variations in substrate availability may account for differences in the occurrence of these evolved enzymes. For example, PETase is secreted from *Ideonella sakaiensis* and other organisms (Yoshida et al., 2021), thereby making direct contact with the PET substrate. On the other hand, MHETase is localized in the periplasmic space and would require internalization of the MHET substrate for the MHET to be utilized as a carbon source (Palm et al., 2019). This difference in accessibility of the two substrates, PET and MHET, may restrict the ability of microbes to evolve MHETase activity. It is also possible that the enzymes that evolved PETase activity were more promiscuous than the tannases that could evolve MHETase activity. Enzyme promiscuity has been shown to confer a fitness advantage in new growth environments (Guzmán et al., 2019). A final possibility could be the toxicity of the final products of MHETase degradation: terephthalic acid and ethylene glycol. Selective pressures could have favored species that do not produce the toxic byproducts of MHETases.

In the present study, we identified *P. stutzeri* as harboring PETase-like and MHETase-like enzymes with high sequence and structural similarity to known plastic degrading enzymes and the ability to degrade polyurethane and PCL. *P. stutzeri* has previously been shown to degrade polyethylene glycol (Obradors & Aguilar, 1991). Interestingly, we noted a difference in the rate of plastic polymer degradation depending on the strain, with ATCC 17832 (Strain designation 419) having the most substantial clearing of both Impranil® and PCL (**Figure 5**). ATCC 17832 was isolated from tartrate-enriched soil in California, US (Rainey et al., 1994) while ATCC 51152 was isolated from drainage water and ATCC 31258 (Strain designation [FERM 2574]) was isolated from soil. It is possible the different isolates possess unique sequence variants of PETase-like or MHETase-like enzymes or in genes from related biochemical pathways that range in degradation capacity. Pseudomonads represent a diverse group of aerobic, gram-negative rods in the class of gammaproteobacteria (Sampedro et al., 2015). They are found in numerous environments, including soil and aquatic ecosystems, and include psychrophilic, halophilic, hydrocarbonoclastic, and heavy metal-tolerant species with diverse metabolic capacity, such as the ability to metabolize multiple types of hydrocarbons (Ghorbannezhad et al., 2022). Future studies are aimed at better understanding and further optimizing degradation capacity from *P. stutzeri* and other candidates. By learning from nature’s solutions, we can continue to develop sustainable solutions for plastic waste management.

## Supplemental materials

**Supplemental Figure 1.**
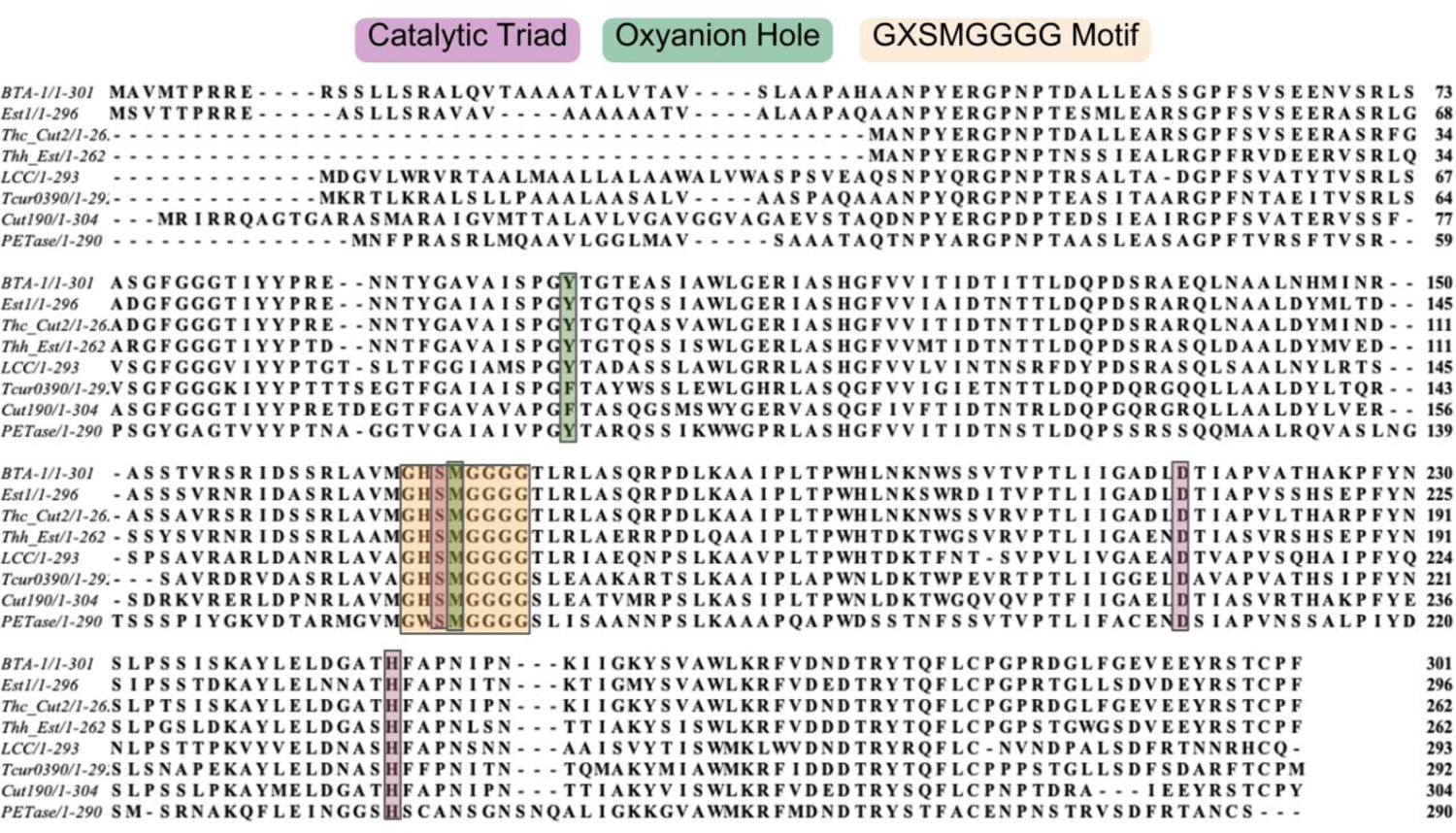

**Supplemental Figure 2.**
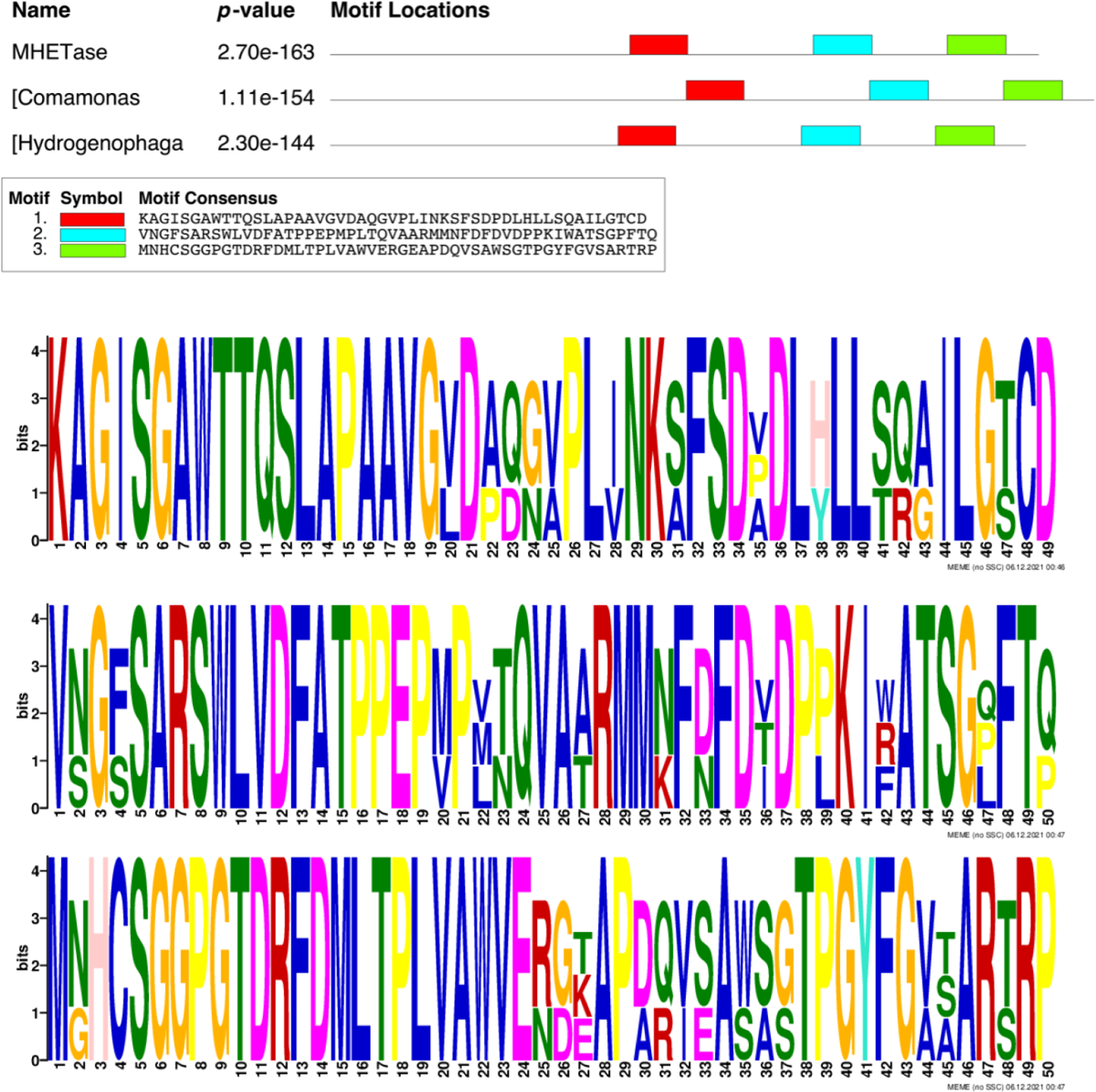

**Supplemental Figure 3.**
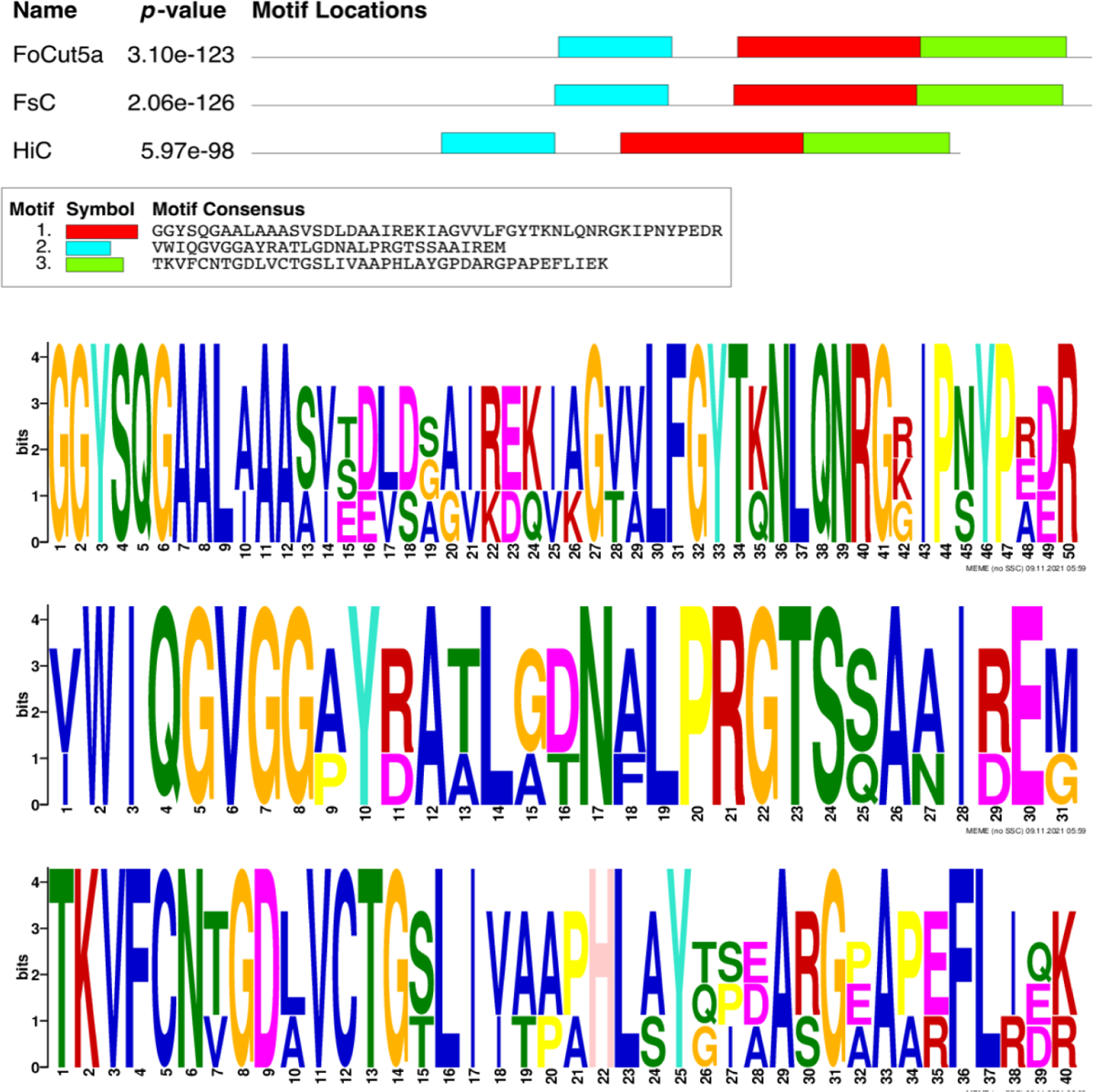

---

**Supplemental Table 1**

https://github.com/alexhong2020/Bioinformatics_Paper

