## Supplemental Figures 1-3 for "A sequence- and structure-based characterization of microbial enzymes identifies P. stutzeri as a plastic-degrading species"

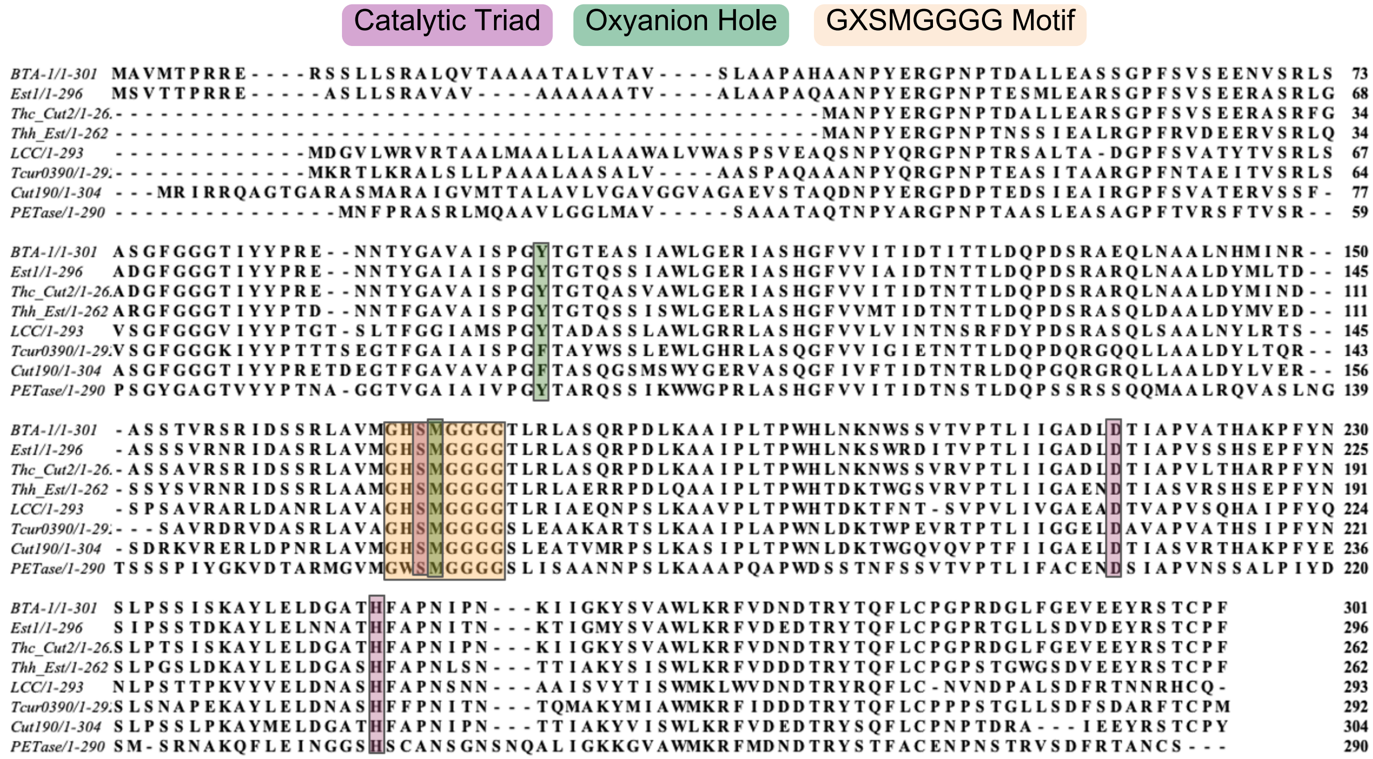

Catalytic Triad
Oxyanion Hole
GXSMGGGG Motif
BTA-1/1-301
Est1/1-296
Thc_Cut2/1-262
Thh_Est/1-262
LCC/1-293
Tcur0390/1-292
Cut190/1-304
PETase/1-290

| Name | *p*-value | Motif Locations |
|---|---|---|
| MHETase | 2.70e-163 | |
| [Comamonas | 1.11e-154 | |
| [Hydrogenophaga | 2.30e-144 | |

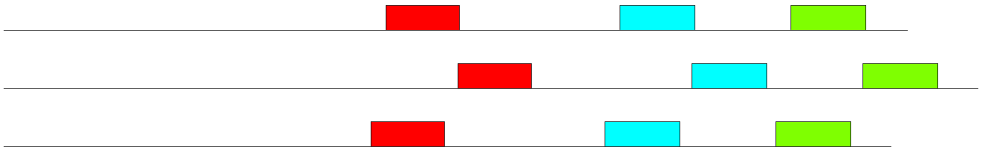

| Motif | Symbol | Motif Consensus |
|---|---|---|
| 1. | | KAGISGAWTTQSLAPAAVGVDAQGVPLINKSFSDPDLHLLSQAILGTCD |
| 2. | | VNGFSARSWLVDFATPPEPMPLTQVAARMMNFDFDVDPPKIWATSGPFTQ |
| 3. | | MNHCSGGPGTDRFDMLTPLVAWVERGEAPDQVSAWSGTPGYFGVSARTRP |

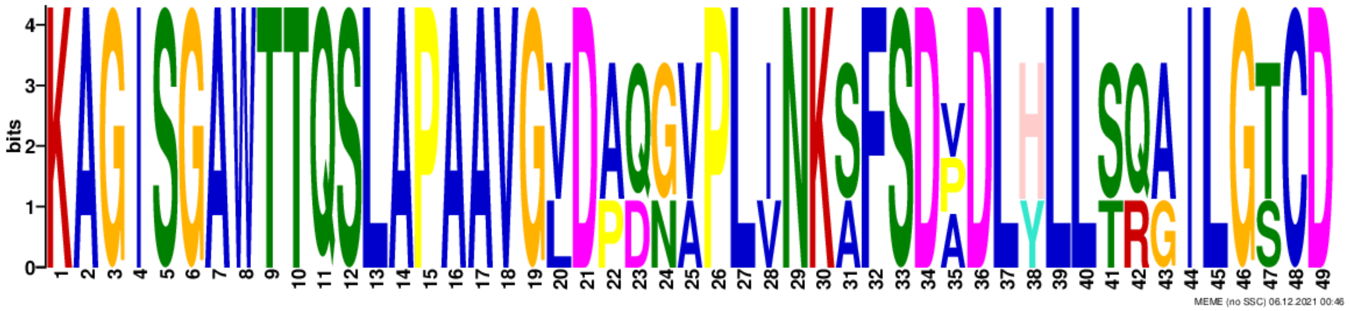


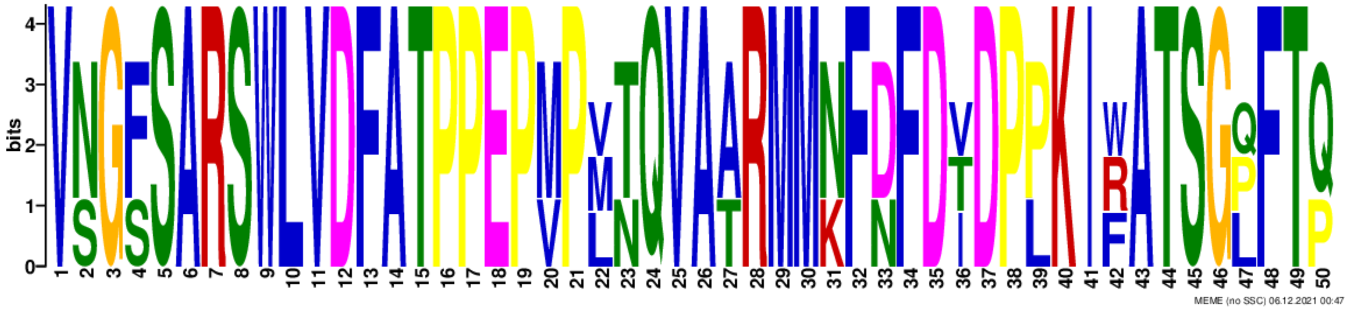


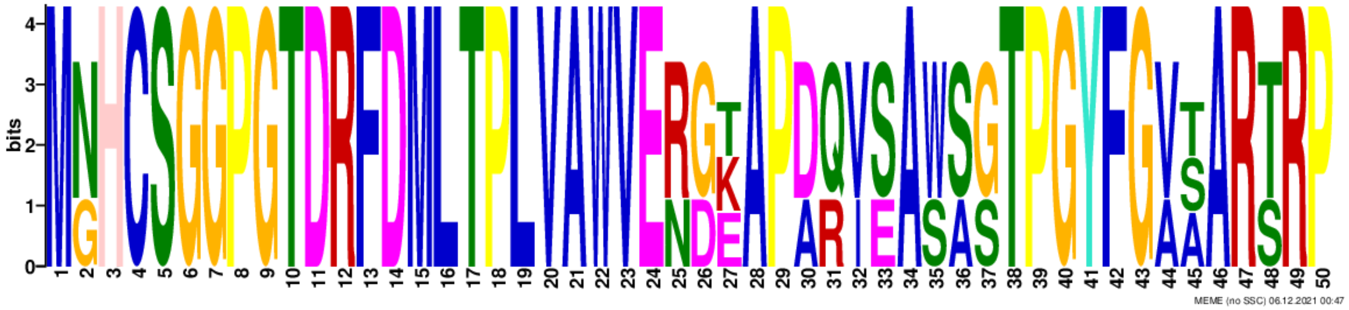

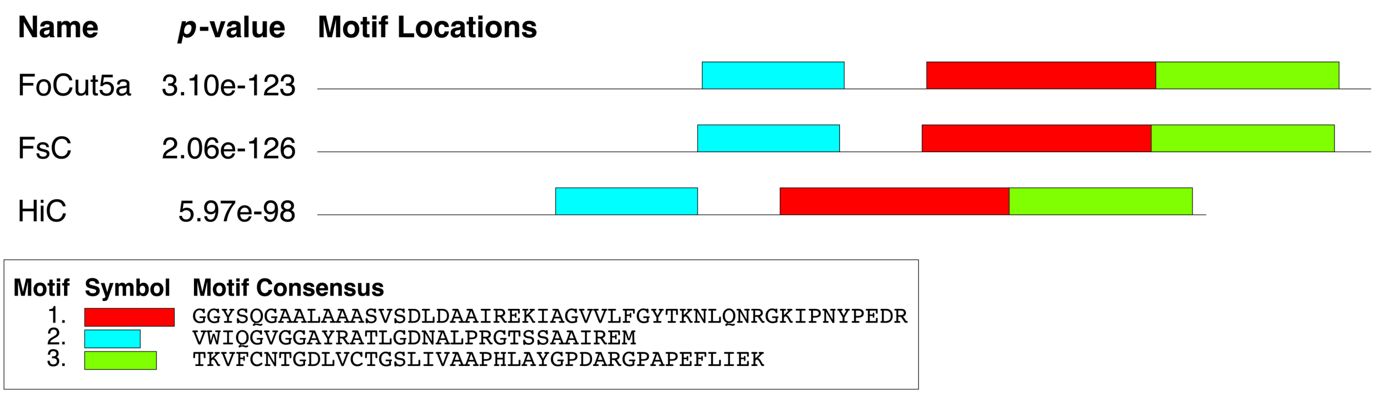
Name
p-value
Motif Locations
FoCut5a
3.10e-123
FsC
2.06e-126
HiC
5.97e-98
Motif
Symbol
Motif Consensus
1. GGYSQGAALAAASVSDLDAAIREKIAGVVLFGYTKNLQNRGKIPNYPEDR
2. VWIQGVGGAYRATLGDNALPRGTSSAAIREM
3. TKVFCNTGDLVCTGSLIVAAPHLAYGPDARGPAPEFLIEK

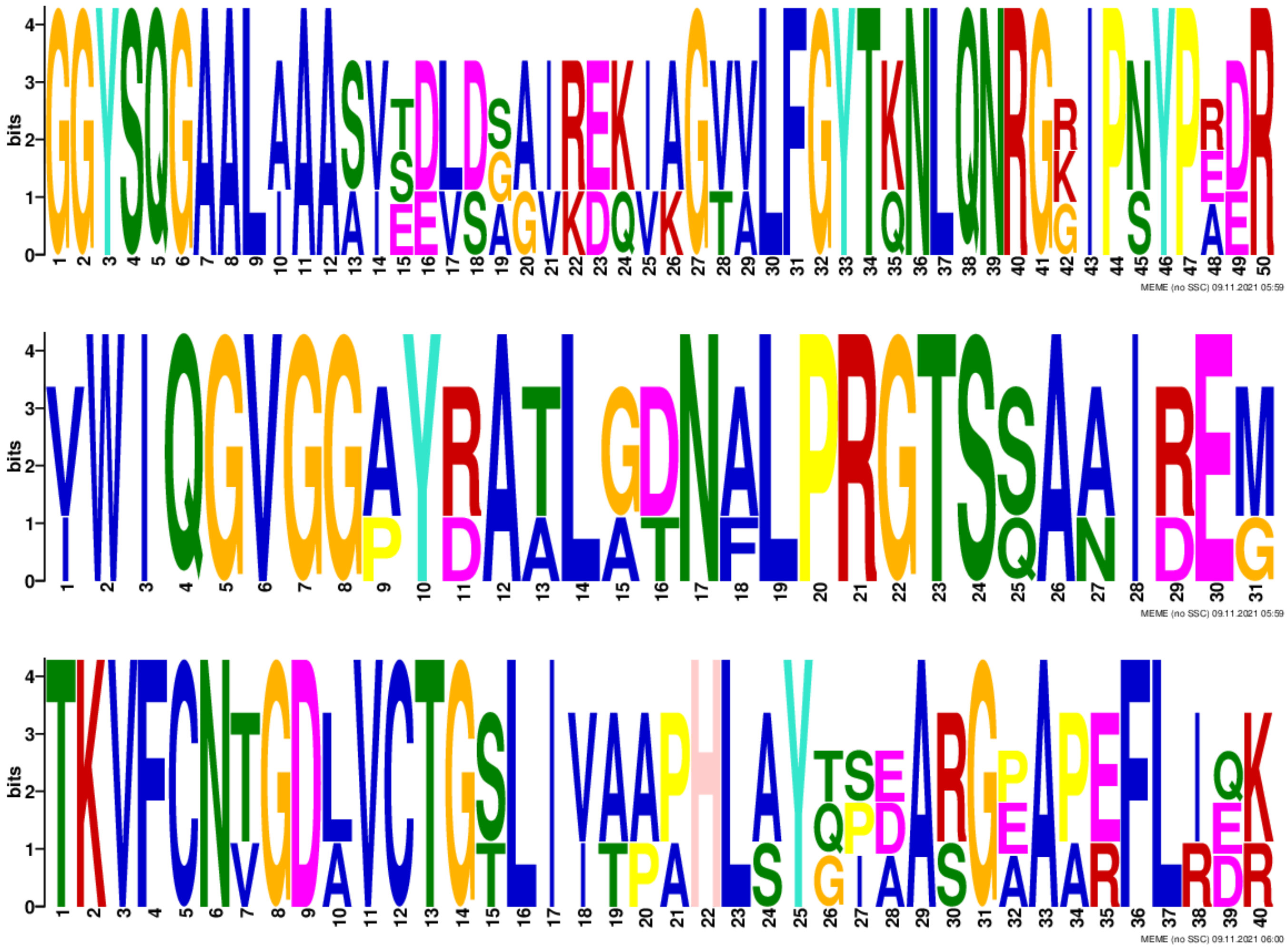
bits
MEME (no SSC) 09.11.2021 05:59
MEME (no SSC) 09.11.2021 05:59
MEME (no SSC) 09.11.2021 06:00
